# Non-canonical Hedgehog signaling regulates spinal cord and muscle regeneration

**DOI:** 10.1101/2020.08.05.238139

**Authors:** Andrew M. Hamilton, Laura N. Borodinsky

## Abstract

Inducing regeneration in injured spinal cord represents one of modern medicine’s greatest challenges. Research from a variety of model organisms indicates that Hedgehog (Hh) signaling may be a useful target to drive regeneration. However, the mechanisms of Hedgehog signaling-mediated tissue regeneration remain unclear. Here we examined Hh signaling during post-amputation tail regeneration in *Xenopus laevis* larvae. We found that while Smoothened (Smo) activity is essential for proper spinal cord and skeletal muscle regeneration, transcriptional activity of the canonical Hh effector Gli is repressed immediately following amputation, and inhibition of Gli1/2 expression or transcriptional activity has minimal effects on regeneration. In contrast, we demonstrate that protein kinase A (PKA) is necessary for regeneration of both muscle and spinal cord, in concert with and independent of Smo respectively, and that its downstream effector CREB is activated in spinal cord following amputation. Our findings indicate that non-canonical mechanisms of Hh signaling are necessary for spinal cord and muscle regeneration.

## Introduction

Injury in low regenerative capacity tissues such as spinal cord represents a persistent challenge in modern medicine, with millions of cases of spinal cord-related disability worldwide (Lee et al., 2014). There are currently no therapies for repairing severe damage in the mammalian central nervous system, but a variety of paradigms have shown great promise in diverse model systems, including stem cell transplantation (Dasari et al., 2014; Lu et al., 2012; Méndez-Olivos et al., 2017), pharmacological and genetic manipulation of cell apoptosis, reactive oxygen species, ion channel function, growth factor and morphogenetic protein signaling and the immune response (Akazawa, 2004; Beck et al., 2003; Fukazawa et al., 2009; Love et al., 2013; Tapia et al., 2017; Tseng et al., 2007, 2010), and direct electrical stimulation (Gomes-Osman et al., 2016). Although diverse cellular processes are involved in neural regeneration, one pathway that has consistently proven to be pro-regenerative is Hedgehog (Hh) signaling. Treatment with Sonic hedgehog (Shh) or with agonists of its main effector, Smothened (Smo), enhances neurological recovery to nerve damage in rats (Bambakidis et al., 2012) possibly by enhancing neural cell proliferation (Bambakidis et al., 2009), while loss of Shh in zebrafish impairs retinal regeneration (Sherpa et al., 2014). Hh signaling is also essential for injury response in a wide variety of tissues, including cardiac muscle (Kawagishi et al., 2018; Singh et al., 2018; Sugimoto et al., 2017), liver (Grzelak et al., 2014), limb (Singh et al., 2012; Yakushiji et al., 2009) and tail (Romero et al., 2018; Taniguchi et al., 2014), making this pathway an attractive therapeutic target.

The Hh pathway is classically known as a primary regulator of organogenesis and tissue homeostasis (Briscoe & Therond, 2013). Hh binding represses its receptor Patched, thereby de-repressing the G-protein coupled receptor Smo. Canonically, Smo activates the transcription factor Gli2, which drives transcription of both Patched and the positive feedback regulator Gli1, as well as a wide variety of genes controlling cell migration, differentiation and especially proliferation (Briscoe & Therond, 2013). In addition to the canonical, Gli-dependent cascade, a number of non-canonical, Gli-independent pathways exist, operating through Src family kinase (Sloan et al., 2015; Yam et al., 2009), the small GTPases Rac1 and RhoA (Ho Wei et al., 2018; Polizio et al., 2011), NF-κB activation via PKC (Qu et al., 2013), and the Ca^2+^-Ampk axis (Teperino et al., 2012), as well as other non-canonical signaling cascades (Belgacem et al., 2016; Teperino et al., 2014). Our own work has shown that Shh drives spontaneous Ca^2+^ spike activity in immature neurons, regulating neuronal differentiation during spinal cord development (Belgacem & Borodinsky, 2011). Moreover, neural tube formation in mouse and frog is associated with a deactivation of canonical Shh signaling (Balaskas et al., 2012; Belgacem & Borodinsky, 2015; Lee et al., 1997), and a concomitant switch to a non-canonical, PKA and Ca^2+^-dependent pathway, which itself contributes to the repression of the canonical, Gli-dependent cascade (Belgacem & Borodinsky, 2015).

To better understand its role in tissue repair, we examined the Hh signaling pathway during tail regeneration in *Xenopus laevis* larvae. We found that Hh signaling is necessary for the regeneration of the spinal cord and skeletal muscle, primarily through Gli-independent pathways, and our results implicate PKA/CREB as an interacting signaling cascade. These findings offer the possibility of enhancing regeneration by differentially targeting canonical and non-canonical Hh pathways.

## Results

### Hedgehog signaling regulates regeneration of muscle and spinal cord

Our previous studies have shown that electrical activity is important for spinal cord and muscle regeneration (Tu & Borodinsky, 2014). In addition, Ca^2+^-dependent activity is necessary for neuronal and muscle cell differentiation during embryonic skeletal muscle and spinal cord development (Borodinsky et al., 2004; Ferrari et al., 1996; Ferrari & Spitzer, 1999; Gu & Spitzer, 1995). This activity drives non-canonical, Ca^2+^-dependent Shh signaling during spinal cord neuron differentiation (Belgacem & Borodinsky, 2011), positioning non-canonical Hh signaling as an excellent candidate for regulating regeneration in muscle and neural tissues. To directly examine the role of Hh in tissue regeneration, the tails of *Xenopus laevis* larvae were amputated and allowed to regenerate in Smo antagonist cyclopamine, Smo agonist SAG, or vehicle control solution, then fixed at 72 h post-amputation (hpa) and stained for mitotic activity (phospho-histone H3, P-H3), spinal cord (Sox2+ neural stem cells (NSCs) lining the spinal cord central canal) and skeletal muscle (12/101+ differentiated skeletal muscle cells) in the regenerate and vicinity (Figure 1A-C).

**Figure 1.**
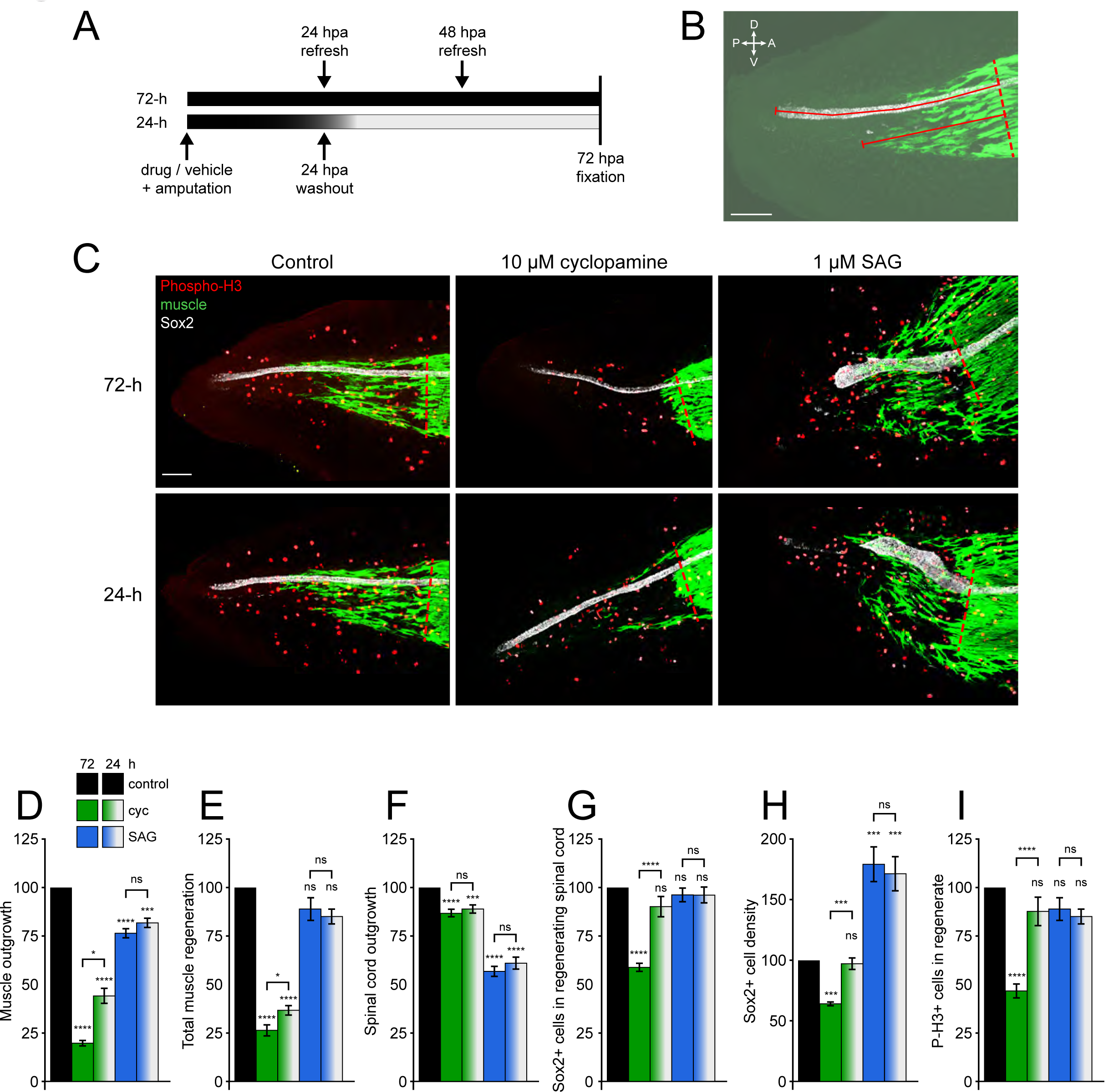
Hedgehog signaling regulates spinal cord and muscle regeneration. Stage 39-40 *Xenopus laevis* larvae were incubated for 24 or 72 h after tail amputation in vehicle (0.1% DMSO, Control), antagonist (10 µM cyclopamine, cyc) or agonist (1 µM SAG) of Smoothened (Smo), and immunostained at 72 h post amputation (hpa). (**A**) Schematic of 24- and 72-h treatments. (**B**) Measurement of outgrowth for regenerated spinal cord and muscle (solid lines) from amputation plane (dashed line). (**C**) Representative z-projections of whole-mount immunostained samples for each group at 72 hpa. Transverse red dashed line indicates amputation plane. Scale bars in (**B**,**C**), 100 µm. (**D**-**H**) Graphs show mean±SEM measurements of regenerated muscle outgrowth (**D**) and total new muscle volume (**E**), regenerated spinal cord outgrowth (**F**), total number (**G**) and per length density (**H**) of Sox2+ cells in the regenerated spinal cord, and number of PH3+ cells (**I**) in the regenerate at 72 hpa as % of cohort-matched control, n of larvae: 25-42 per group, N of experiments≥3. *p<0.05, ***p<0.001, ****p<0.0001, ns: not significant, ordinary one-way ANOVA, Brown-Forsythe and Welch ANOVA, or Kruskal-Wallis test, followed by Tukey’s, Dunnett’s T3 or Dunn’s multiple comparisons test, respectively, according to prior normality and equality of SDs tests within and between groups.

Smo modulation alters both muscle and spinal cord regeneration. Inhibition of Smo for 72 hpa with cyclopamine reduces muscle outgrowth into the regenerating tail (Figure 1D) and total regenerated muscle volume (Figure 1E). Cyclopamine also impairs spinal cord outgrowth (Figure 1F), and reduces the total number of NSCs in the newly regenerated spinal cord (Figure 1G), resulting in new spinal cord showing a lower NSC density than in control larvae (Fig. 1H), which suggests that inhibiting Smo interferes with activation/proliferation of NSCs. Enhancing Smo activity with SAG, however, has no effect on total muscle regeneration (Figure 1E), and even shows a modest inhibitory effect on muscle outgrowth (Figure 1D). In addition, SAG treatment results in disorganized muscle regeneration and ectopic outgrowth of new muscle fibers dorsally and ventrally from intact axial musculature in the tail stump (Figure 1C), indicating dysregulated muscle morphogenesis. SAG also reduces the outgrowth of the spinal cord (Figure 1F) without altering the total number of new NSCs (Figure 1G), resulting in a truncated, wide spinal cord with higher NSC density than in control larvae (Figure 1C,H). Similarly, overall mitotic activity at 72 hpa is not affected by SAG but is reduced by cyclopamine (Figure 1I). These results argue that Smo activity is essential for NSC proliferation, and that ectopically elevated Smo signaling interferes with the normal progression from NSC proliferation to spinal cord outgrowth.

Overall, these findings are consistent with previous studies demonstrating a role for Hh signaling in cell proliferation (Amankulor et al., 2009; Lai et al., 2003; Romero et al., 2018; Rowitch et al., 1999), differentiation (Belgacem & Borodinsky, 2011) and migration (Armstrong et al., 2017; Gordon et al., 2018), and demonstrate that balanced Hh signaling is necessary for proper tissue regeneration.

In addition, we examined the interval during which Hh signaling is necessary for regeneration by comparing 72-h exposure with treatment for only the first 24 h of 72 total (Figure 1A; 24-h vs 72-h). We found that the effects of 24-h exposure to SAG were comparable to the full 72-h treatment for all metrics of both muscle (Figure 1D,E) and spinal cord (Figure 1F,G,H) regeneration, as well as overall cell proliferation in the regenerate (Figure 1I), and gave the same ectopic muscle overgrowth phenotype and shortened spinal cord (Figure 1C). Cyclopamine treatment for only 24-h, however, showed weaker effects than the full 72-h; the number and density of NSCs in the newly formed spinal cord, as well as the number of mitotic cells in the regenerate are unchanged by 24-h cyclopamine (Figure 1G,H,I), while reductions in muscle outgrowth and volume are smaller than with 72-h cyclopamine incubation (Figure 1D,E). In contrast, the reduction in spinal cord outgrowth is comparable between 24 and 72-h cyclopamine treatments (Figure 1F).

These results suggest that endogenous Hh signaling plays an early and ongoing role in maintaining the regenerative proliferation necessary for tissue growth, and indicate that balanced Hh signaling is necessary for spinal cord and muscle regeneration. Furthermore, the persistent effects of Hh signaling enhancement during the first 24 hours indicate an early critical period during which balanced Hh activity is necessary for new tissue morphogenesis.

### Canonical hedgehog signaling is rapidly repressed following tail amputation

Next, we examine the endogenous activity of canonical Hh signaling following amputation using a dual-cassette transcriptional activity reporter plasmid with simultaneous Gli1/2-dependent expression of enhanced green fluorescent protein (EGFP), and constitutive expression of near-infrared fluorescent protein as a normalizing factor for reporter expression (iRFP670; Gli reporter; Figure 2A). When injected into 2-4-cell-stage embryos, this construct is distributed in a mosaic fashion throughout the larva, giving an expression-normalized readout of Gli1/2 transcriptional activity (EGFP:iRFP670 ratio, Figure 2A’). We validated this construct by treating Gli reporter-injected embryos with SAG to increase EGFP:iRFP670 signal during neural plate development, a period known to exhibit high canonical Hedgehog signaling activity (Lee et al., 1997; Belgacem and Borodinsky, 2015) (Figure 2-figure supplement 1).

**Figure 2.**
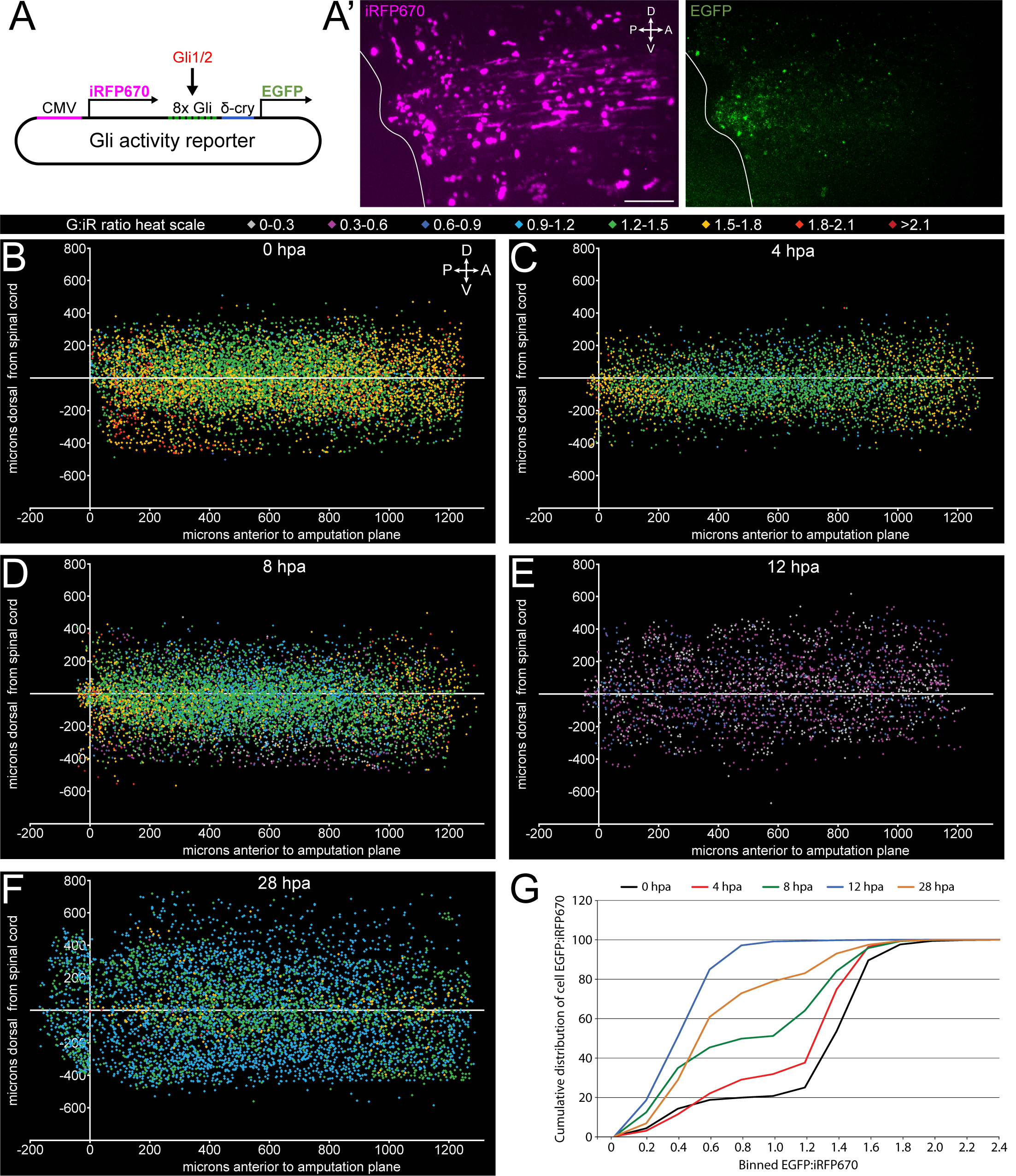
Canonical, Gli-dependent Hedgehog signaling is downregulated following tail amputation. Tails of stage 39-40 larvae expressing the Gli transcriptional activity reporter were amputated and imaged live at intervals from 0-28 hpa. (**A**) Schematic of bicistronic reporter plasmid. Constitutive promoter CMV drives expression of iRFP670, and minimal δ-crystallin promoter juxtaposed to 8 Gli-binding sites drives expression of EGFP. (**A’**) Representative z-projections of normalizing factor (iRFP670) and Gli transcriptional activity reporter (EGFP) in amputated larval tail at 12 hpa. White outline represents the edge of the tail. Scale bar, 100 µm. (**B**-**F**) 2D composite of EGFP:iRFP670 (G:iR ratio) intensity displayed by heat scale at indicated hpa from combined iRFP670+ cells. Cell location displayed by position relative to amputation plane(x) and dorsal or ventral to the spinal cord (y). (**G**) Cumulative distribution of EGFP:iRFP670 ratios by timepoint. All cumulative distribution curves significantly different from control (0 hpa); n of larvae ≥ 12; p<0.001, all time points compared to 0 hpa, Kolmogorov-Smirnov post-hoc test. The online version of this article includes the following figure supplement for figure 2: **Figure supplement 1**. Positive control for Gli transcriptional activity reporter. Neural plate stage embryos (18 hpf) expressing Gli transcriptional activity reporter were incubated in the presence or absence of 10 nM SAG for 5 h followed by live confocal imaging with 488 and 647 nm lasers to acquire the signal corresponding to Gli transcriptional activity (EGFP) and expression level for the reporter (iRF670). Individual cells displayed by iRFP670 (x) vs EGFP (y) fluorescence intensities in control and experimental embryos. 9 embryos per condition.

When each iRFP670+ cell is assigned a heat scale color by EGFP:iRFP670 intensity, and displayed in two-dimensional space by its position relative to the spinal cord (Y) and the amputation plane (X) we find that relative to 0 hpa (Figure 2B), by 4 hpa there is a broad reduction in Gli-reporter signal which appears 300-800 μm anterior to the amputation plane (Figure 2C). This decrease expands and deepens through 8 hpa, until reaching a minimum at 12 hpa, before showing partial recovery at 28 hpa (Figure 2D-F). When the cumulative distribution of EGFP:iRFP670 ratios is compared by timepoint, we find significant reductions in the proportion of higher Gli-activity cells at all timepoints post amputation (Figure 2G).

These data indicate that amputation rapidly induces widespread inhibition of canonical Gli-dependent signaling.

### Hedgehog-dependent spinal cord and muscle regeneration are primarily non-canonical

Our data suggest that while Smo-dependent Hh signaling is necessary for proper spinal cord and muscle regeneration, canonical, Gli-dependent activity is downregulated immediately following tail amputation; therefore, we directly addressed the necessity of Gli1/2 activity during tail regeneration. GANT61 is a small molecule inhibitor of the transcriptional activity of Gli1 and Gli2, the primary activators of downstream canonical Hh signaling (Lauth et al., 2007). Using our Gli reporter, we demonstrate that treatment with 10 µM GANT61 reduces Gli transcriptional activity in *X*. *laevis* larvae (Figure 3-figure supplement 1).

**Figure 3.**
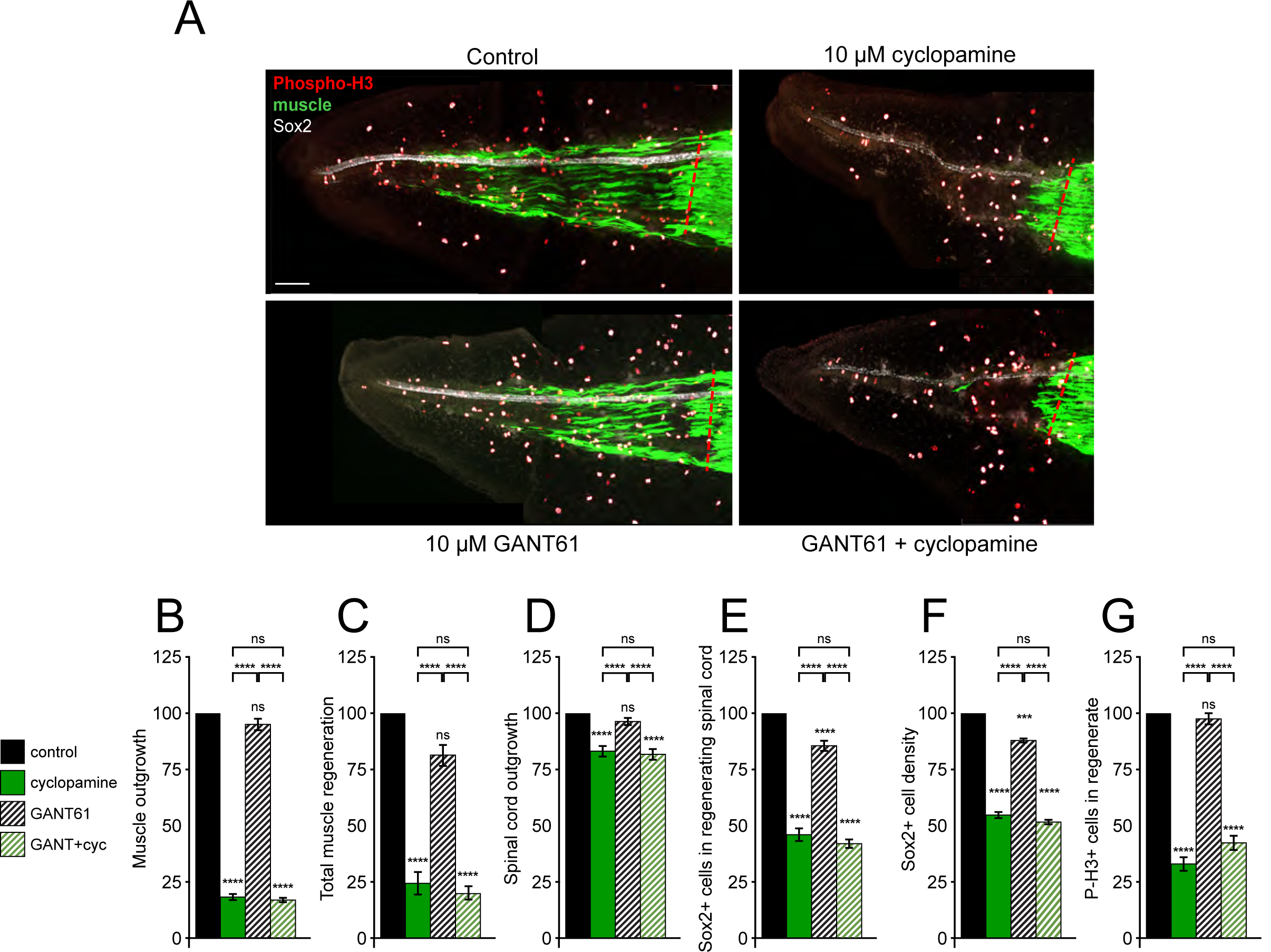
Gli1/2 transcriptional activity is not necessary for spinal cord and muscle regeneration. Stage 39 larvae were incubated for 72 h after tail amputation in vehicle (0.1% DMSO, Control) or Gli1/2 antagonist (10 µM GANT61, GANT) and/or 10 µM cyclopamine (cyc), then whole mount immunostained. (**A**) Images show representative z-projections for each group at 72 hpa. Transverse red dashed line indicates amputation plane. Scale bar, 100 µm. (**B**-**F**) Graphs show mean±SEM measurements of regenerated muscle outgrowth (**B**) and total volume (**C**), regenerated spinal cord outgrowth (**D**), total number (**E**) and per length density (**F**) of Sox2+ cells in the regenerated spinal cord, and overall number of PH3+ cells (**G**) in the regenerate at 72 hpa as % of cohort-matched control, n of larvae: 13-31 per group, N of experiments≥3. ***p<0.001, ****p<0.0001, ns: not significant, ordinary one-way ANOVA, Brown-Forsythe and Welch ANOVA, or Kruskal-Wallis test, followed by Tukey’s, Dunnett’s T3 or Dunn’s multiple comparisons test, respectively, according to prior normality and equality of SDs within and between groups. The online version of this article includes the following figure supplement for figure 3: **Figure supplement 1**. Treatment with GANT61 inhibits Gli1/2 transcriptional activity. Stage 39 larvae expressing the Gli transcriptional activity reporter were incubated with 10 µM GANT61 and confocally live-imaged with 488 and 647 nm lasers to acquire the signal corresponding to Gli transcriptional activity (EGFP) and expression level for the reporter (iRF670), at 0 (**A**), 6 (**B**) and 18 h (**C**) of incubation. Shown are paired EGFP and iRFP fluorescence intensities for individual cells at the indicated time points. N of larvae: 8.

Treatment of tail-amputated larvae with GANT61 for 72 h (Figure 3A) results in only a modest reduction in the number of NSCs (Figure 3E), reducing Sox2 density in the regenerating spinal cord (Figure 3F), but does not significantly alter any other regeneration or proliferation metrics in either muscle or spinal cord (Figure 3B-D,G). This contrasts sharply with cohort-paired cyclopamine treatment, which shows reductions in all regeneration metrics for muscle and spinal cord (Figure 3). Furthermore, when GANT61 treatment is combined with cyclopamine, the results are indistinguishable from cyclopamine treatment alone for all regeneration metrics (Figure 3).

In addition to pharmacological inhibition of Gli1/2 activity, we used a translation-blocking morpholino to downregulate protein abundance of Gli2, the primary activator of canonical Hh signaling. As Gli2 signaling is indispensable for early embryogenesis, we coinjected a complementary, UV photolabile blocker morpholino (Photo-MO) which binds to and disables the matched Gli2 morpholino (Gli2-MO), allowing UV-dependent uncaging of Gli2-MO at later developmental periods (Figure 4A). Exposure of uninjected embryos to UV does not reduce any regeneration metrics (Figure 4-figure supplement 1A-D), while activation of Gli2-MO has no significant effect on regeneration of muscle or spinal cord, with the exception of a small reduction in spinal cord outgrowth (Figure 4B-G), despite notable downregulation of full length Gli2 protein levels (Figure 4-figure supplement 1E).

**Figure 4.**
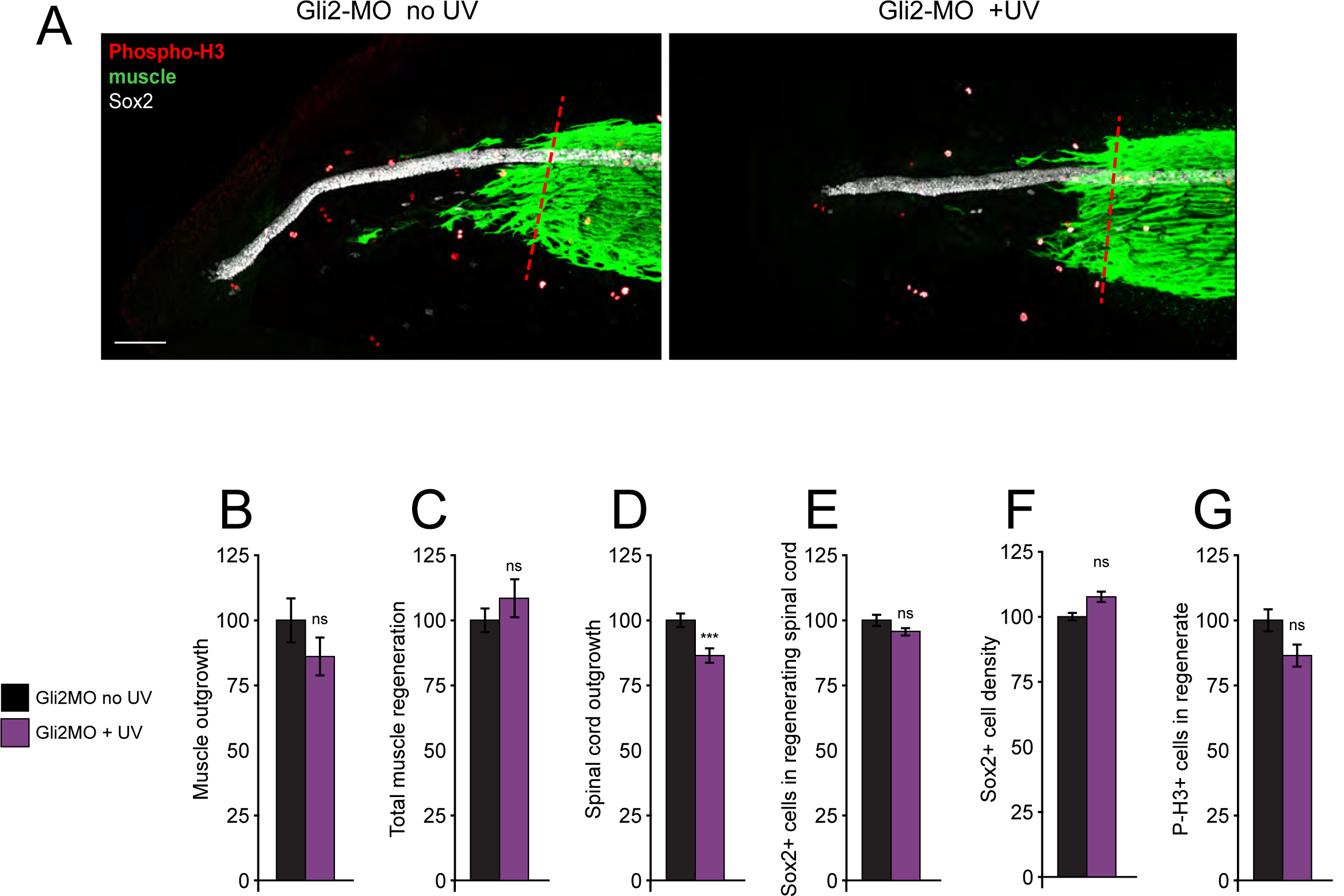
Genetic downregulation of canonical Hh-Smo signaling does not affect spinal cord and muscle regeneration. Larvae containing Gli2 morpholino (Gli2-MO) bound to photo-morpholino were UV illuminated (+UV) or not (control, no UV) for 30 min, 28 h before (stage 28) amputation (stage 39-40), to uncage morpholino and induce downregulation of Gli2 expression. (**A**) Images show representative z-projections of samples for each group at 72 hpa. Transverse red dashed line indicates amputation plane. Scale bar, 100 µm. (**B**-**F**) Graphs show mean±SEM measurements of regenerated muscle outgrowth (**B**) and total volume (**C**), regenerated spinal cord outgrowth (**D**), total number (**E**) and per length density (**F**) of Sox2+ cells in the regenerated spinal cord, and overall number of PH3+ cells (**G**) in the regenerate at 72 hpa as % of cohort-matched control, n of larvae: 16-35 per group, N of experiments≥3. ***p<0.001, ns: not significant, unpaired t-test, Welch’s t-test, or Kolmogorov-Smirnov test, according to prior normality and equality of SDs tests within and between groups. The online version of this article includes the following figure supplement for figure 4: **Figure Supplement 1**. UV treatment does not reduce regeneration, and uncages Gli2-MO. Uninjected larvae (**A**-**D**) or larvae containing Gli2-MO bound to photo-morpholino (**E**) were UV illuminated (+UV) or not (control, no UV) for 30 min, 28 h (stage 28) before amputation (stage 39-40). Amputated samples were processed at 72 hpa for whole-mount immunostaining (**A**-**D**) or frozen immediately for Western blot assays (**E**). (**A**-**D**) Graphs show mean±SEM measurements of regenerated muscle outgrowth (**A**) and total volume (**B**), regenerated spinal cord outgrowth (**C**) and overall number of PH3+ cells (**D**) in the regenerate at 72 hpa as % compared to control, n of larvae: 12-19 per group. *p<0.05, ns: not significant, unpaired t-test. (**E**) Representative Western blot assay. Predicted full-length Gli2 MW≥168 kD indicated by arrow. Lamin B1 (bottom) was used as loading control for nuclear fraction.

These findings indicate that canonical Hh signaling is, for the most part, not necessary for spinal cord and muscle regeneration, and suggest that Smo-dependent regeneration recruits a non-canonical signaling pathway.

### PKA signaling is necessary for regeneration

In previous studies, we found that Ca^2+^ activity is associated with both muscle regeneration (Tu and Borodinsky, 2014) and PKA-dependent non-canonical Shh signaling through Smo during embryonic spinal cord development (Belgacem & Borodinsky, 2011, 2015). To examine a potential role for PKA activity during tail regeneration, we treated amputated larvae with the specific PKA inhibitor KT5720 (Figure 5A). Inhibiting PKA diminishes all regeneration metrics, reducing outgrowth of muscle (Figure 5B) and spinal cord (Figure 5D), total muscle regeneration (Figure 5C), and spinal cord NSC (Figure 5E) and mitotic cell counts (Figure 5G). The reduction in NSC number when PKA is inhibited is approximately proportional to the reduction in spinal cord outgrowth, resulting in only a slight increase in NSC density (Figure 5F), despite the fact that PKA inhibition leads to a larger overall decrease in spinal cord regeneration than when inhibiting Smo (Figure 5D). This suggests that PKA participates in spinal cord regeneration beyond the initial activation and proliferation of NSCs, while Smo signaling is particularly important for NSC proliferation.

**Figure 5.**
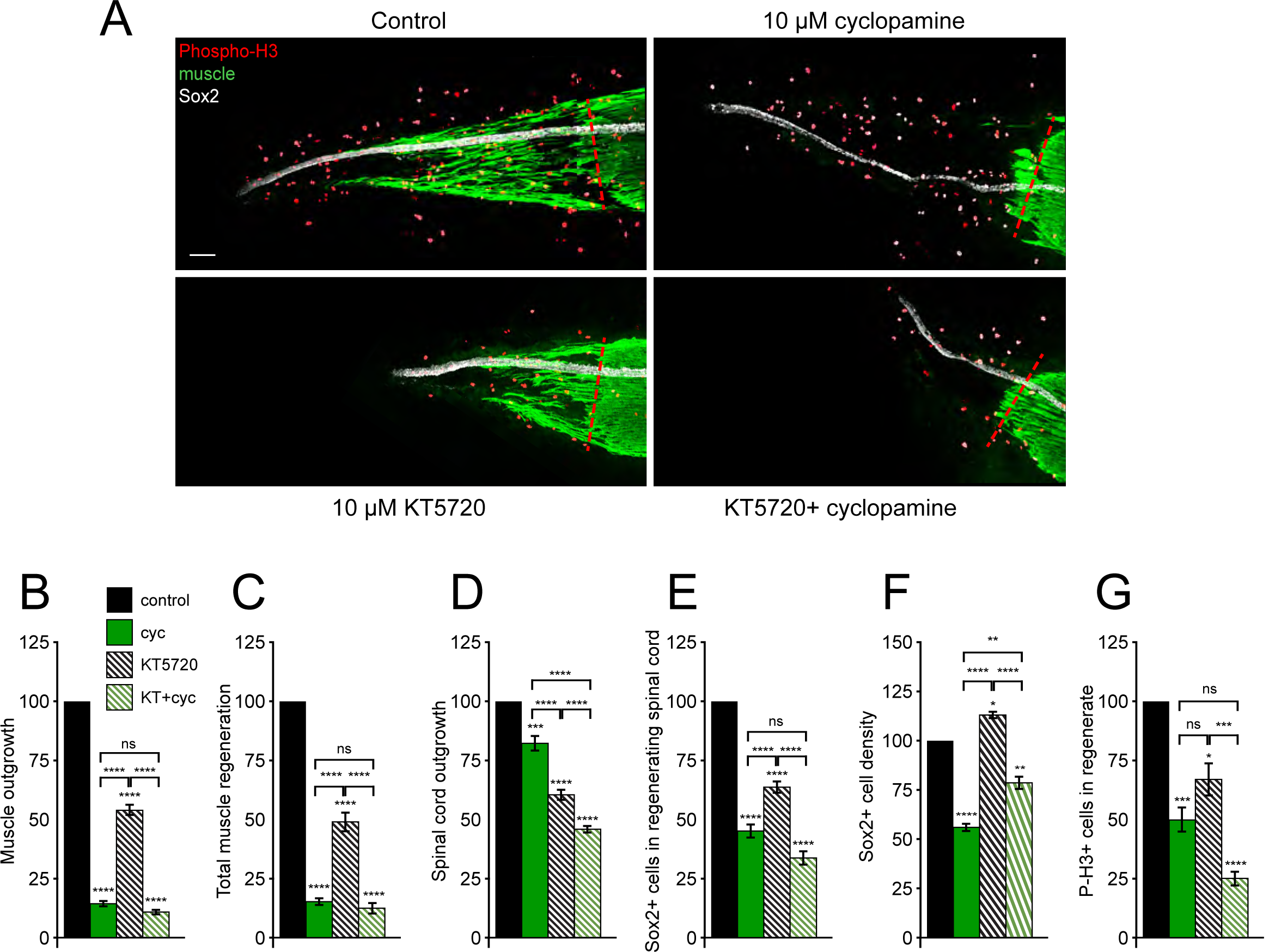
Tissue-specific interaction between PKA and Smo signaling in regulation of spinal cord and muscle regeneration. Stage 39 larvae were incubated for 72 h after tail amputation in vehicle (0.1% DMSO, Control), PKA antagonist (10 µM KT5720, KT) or/and 10 µM cyclopamine (cyc). (**A**) Images show representative samples for each group at 72 hpa. Transverse red dashed line indicates amputation plane. Scale bar, 100 µm. (**B**-**F**) Graphs show mean±SEM measurements of regenerated muscle outgrowth (**B**) and total volume (**C**), regenerated spinal cord outgrowth (**D**), total number (**E**) and per length density (**F**) of Sox2+ cells in the regenerated spinal cord, and overall number of PH3+ cells (**G**) in the regenerate at 72 hpa as % of cohort-matched control, n of larvae: 11-29 per group, N of experiments≥3. *p<0.05, **p<0.01, ***p<0.001, ****p<0.0001, ns: not significant, ordinary one-way ANOVA, Brown-Forsythe and Welch ANOVA, or Kruskal- Wallis test, followed by Tukey’s, Dunnett’s T3 or Dunn’s multiple comparisons test, respectively, according to prior normality and equality of SDs within and between groups.

Simultaneous treatment with KT5720 and cyclopamine reveals tissue-specific epistasis: inhibition of Smo is epistatic over PKA inhibition for muscle outgrowth (Figure 5B) and total new muscle formation (Figure 5C), suggesting that PKA acts upon muscle regeneration downstream of Smo. In contrast, dual KT5720/cyclopamine treatment results in additive inhibition of spinal cord outgrowth (Figure 5D), but no significant additivity in reduction of NSC (Figure 5E) or mitotic cell counts (Figure 5G). Interestingly, simultaneous inhibition of PKA and Smo slightly ameliorates the reduction in NSC density observed with cyclopamine alone (Figure 5F), reinforcing the concept that PKA acts upon spinal cord regeneration in a mechanistically distinct manner from Smo. This suggests that PKA and Smo act on spinal cord regeneration in a parallel but coordinated manner.

Since PKA is known to be a potent regulator of a wide variety of signaling pathways, we directly examined one of PKA’s primary downstream effectors, the transcription factor CREB, which is activated by non-canonical Hh signaling in the embryonic spinal cord (Belgacem and Borodinsky, 2015). To examine endogenous CREB activity following amputation, we fixed larvae at intervals from 0-48 hpa and stained for activated phospho-Ser133-CREB (P-CREB, (Belgacem & Borodinsky, 2015; Gonzalez & Montminy, 1989)). We found that P-CREB signal is especially strong in the skin around the regenerating tail tip, making isolation of P-CREB signal in muscle unfeasible (Figure 6-figure supplement 1). However, we were able to isolate P-CREB staining from the Sox2-labeled region of the spinal cord (Figure 6A). We find that the density of P-CREB+ cells in the spinal cord within the amputated tail stump remains constant over 48 hpa, comparable to pre-amputation controls (Figure 6B). Within the regenerating spinal cord, however, we observe a significant increase in P-CREB+ cells at 4 hpa, compared to the intact stump (Figure 6B). This increase in P-CREB+ cell density progressively diminishes at 12 and 24 hpa and is no longer enriched at any region in the regenerating spinal cord by 48 hpa (Figure 6B). If we further separate total P-CREB+ cells into dim and bright P-CREB intensity groups, we see that the increase in P-CREB+ cells at 4 hpa is largely due to an increase in bright cells which is greatly reduced by 12 hpa and no longer apparent by 24 hpa, indicating temporal regulation of CREB activation at the amputation site within the newly injured spinal cord (Figure 6D). Additionally, we find that by 24 hpa, the regenerating tail is largely populated by dim P-CREB+ cells, while the intact stump shows a distinctive enrichment in bright P-CREB+ cells at 48 hpa (Figure 6C, D). This suggests that CREB activity is differentially regulated spatially and temporally in the spinal cord, with a rapid, transient increase in CREB activation in the regenerate proximal to the amputation site, followed by delayed activation in the intact stump.

**Figure 6.**
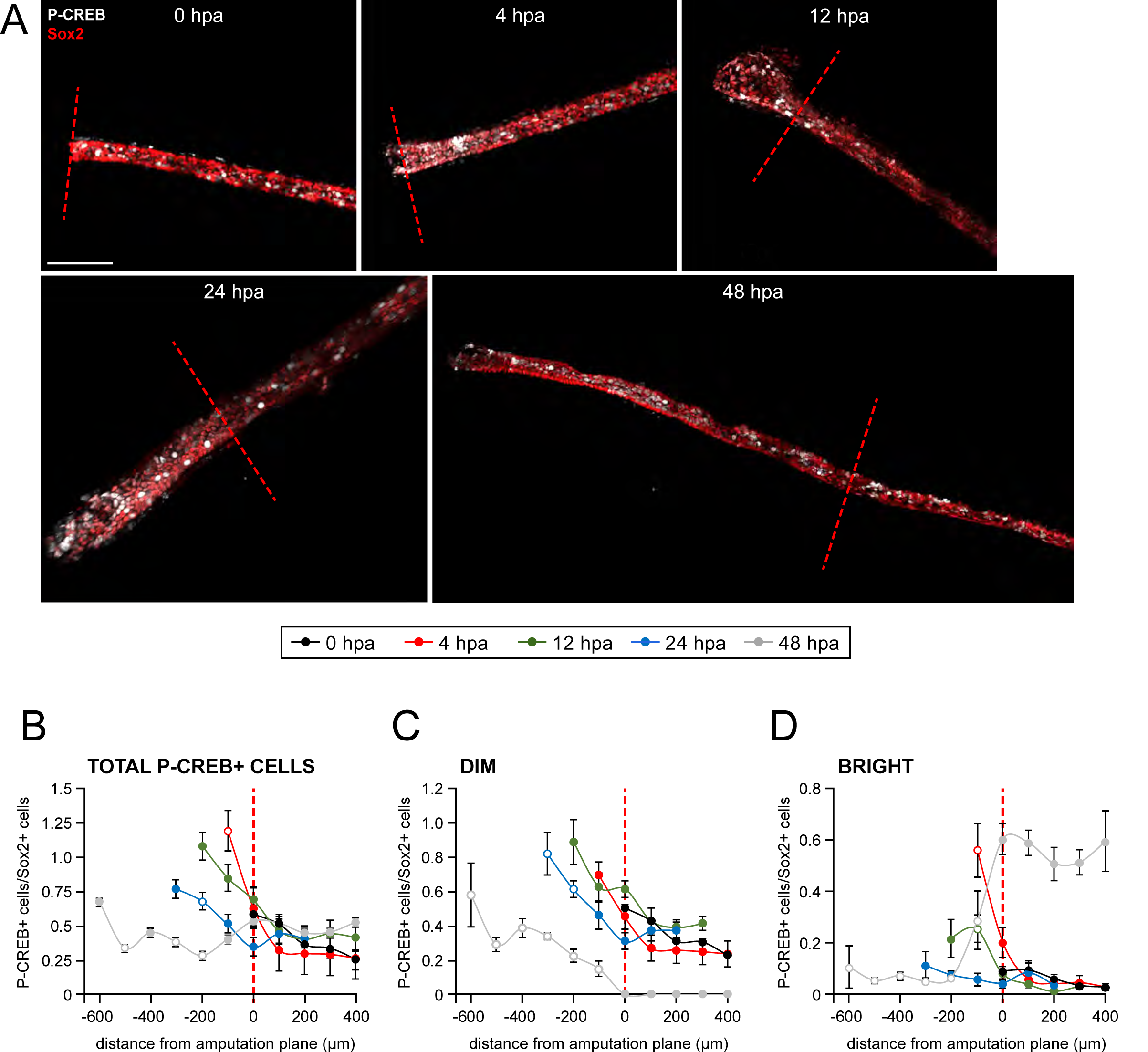
Spatiotemporal activation of CREB in the injured and regenerated spinal cord. Stage 39- 40 larvae were amputated and processed for whole-mount immunostaining at the indicated hpa, with the exception of the 0 hpa group, which was first fixed and then amputated to represent the pre-amputation group. (**A**) Images show representative z-projections of immunostained samples for each group that were digitally processed to isolate spinal-cord associated P-CREB. Transverse red dashed line indicates amputation plane. Scale bar, 100 µm. (**B**-**D**) Graphs show mean±SEM number of total (**B**), dim (**C**) and bright (**D**) P-CREB+ cells normalized to number of Sox2+ cells in 100-µm sections of spinal cord anterior (positive) and posterior (negative) to the amputation plane (0). N of larvae≥5 per group. Open circles denote p<0.05 vs time-matched amputation plane region (0 µm), 1-way ANOVA. The online version of this article includes the following figure supplement for figure 6: **Figure Supplement 1**. Unedited images from samples featured in Figure 6. Stage 39 larvae were amputated and processed for whole-mount immunostaining at the indicated hpa, with the exception of the 0 hpa group, which was first fixed and then amputated to represent the pre-amputation group. Images show representative total z-projections of whole-mount immunostained samples for each group corresponding to the edited images shown in Figure 6. Transverse red dashed line indicates amputation plane. Scale bar, 100 µm.

## Discussion

Our work demonstrates that Hh signaling is necessary for regeneration of muscle and spinal cord in *Xenopus* larvae. These findings are in keeping with previous discoveries showing Hh signaling involvement in the regeneration of a variety of tissues including liver, heart and limb (Grzelak et al., 2014; Singh et al., 2012, 2018), in addition to *Xenopus* larval tail (Taniguchi et al., 2014), as well as the fact that Hh pathway activation is enhanced at the site of tail amputation in *X*. *laevis* (Taniguchi et al., 2014) and zebrafish (Romero et al., 2018).

Under control conditions we observe a close correlation between muscle and spinal cord outgrowth, when examined on a per sample basis. This correlation is decoupled by enhancing Smo signaling, which allows muscle outgrowth in the absence of normal spinal cord outgrowth, and by inhibiting it, which blocks muscle regeneration almost entirely (Figure 7A). In contrast, PKA inhibition reduces all regeneration metrics without altering the interrelationship between the extent of regeneration of both tissues, and inhibiting Gli1/2 has no discernable effect on correlated outgrowth (Figure 7A). This suggests that balanced non-canonical Hh signaling is necessary for coordinated regeneration of spinal cord and muscle (Figure 7B).

**Figure 7.**
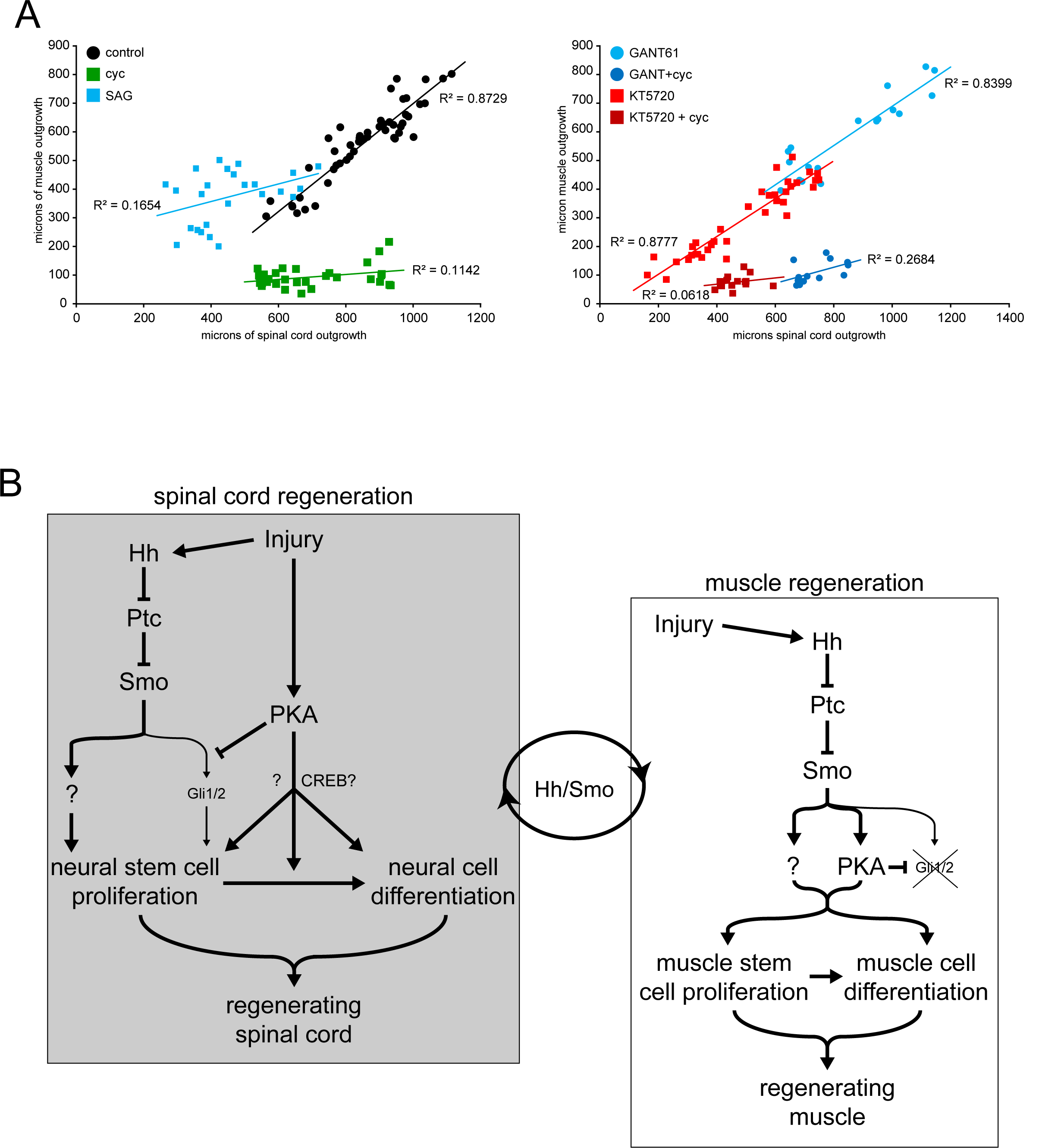
Coordination of spinal cord and muscle regeneration by non-canonical Hh-Smo and PKA signaling pathways. (**A**) Correlation between spinal cord outgrowth and muscle outgrowth. Individual samples for each treatment are displayed by total outgrowth for spinal cord (X) and muscle (Y). Simple linear regression lines are fit for each treatment, and R2 goodness of fit values displayed by each line. N=13-39 samples per group. (**B**) Model for Hh-dependent regulation of spinal cord and muscle regeneration. Injury recruits non-canonical Hh signaling in the spinal cord to activate neural stem cells for replenishing of the regenerating spinal cord. Injury also activates PKA that acts independently from Smo action on neural stem cell proliferation to promote spinal cord regeneration. Non-canonical Hh signaling is also essential for muscle regeneration starting from the initial stages post injury, where PKA appears to be downstream of Smo activation. The apparent coordination between the magnitude of regeneration in spinal cord and muscle is dependent on non-canonical Hh/Smo signaling.

The regenerated spinal cord under Smo inhibition exhibits around half of the number of NSCs by 72 hpa compared to control animals. This suggests that with regards to spinal cord regeneration, Hh-Smo signaling is particularly important for NSC proliferation. Moreover, pharmacological enhancement of Smo signaling leads to simultaneous NSC proliferation and complete blockade of spinal cord morphogenesis, further supporting the model of Smo signaling acting predominantly upon NSC activation and proliferation upon injury (Figure 7B).

Smo signaling is also essential for muscle regeneration and arguably for the initial stages of muscle stem cell activation and proliferation, since inhibiting Smo signaling during the first 24 hpa is enough to strongly decrease the replenishing of skeletal muscle (Figure 7B). Moreover, it appears that the first 24 h post injury are a critical period for Smo-dependent muscle stem cell activation, unlike neural stem cells, which seem to be able to be activated and proliferate even after an initial 24-h of Smo inhibition. It remains to be determined whether Smo-dependent muscle regeneration is acting directly on muscle stem cells or indirectly through signals from the spinal cord. Nevertheless, the evidence that in the presence of overactive Smo, both the spinal cord and muscle exhibit aberrant outgrowth, along with published work from others demonstrating the necessity of the spinal cord for proper tail regeneration (Taniguchi et al., 2008), support a potential interaction between Smo-mediated spinal cord and muscle regeneration (Figure 7A,B).

Additionally, we demonstrate that canonical, Gli-dependent Hh signaling is endogenously downregulated after tail amputation, and further inhibiting this pathway either pharmacologically or genetically only marginally affects muscle and spinal cord regeneration, suggesting that Hh-Smo is acting primarily through a Gli-independent pathway (Figure 7B). In contrast, CREB activity is recruited immediately after injury, at the amputation site and within the regenerating spinal cord, and fades within the first 24 h of spinal cord regeneration. Moreover, PKA activity, unlike Gli, is necessary for efficient regeneration of both muscle and spinal cord, although it is possible that PKA inhibition impairs muscle regeneration by reducing spinal cord regeneration, as KT5720 does not alter the close correlation between the outgrowth of spinal cord and muscle (Figure 7A). This distinct recruitment of a non-canonical Hh signaling pathway resembles the switch in Hh signaling observed during embryonic stages of early spinal cord development, when canonical, Gli-activating Shh signaling is restricted to the early neural plate stages (Balaskas et al., 2012; Belgacem & Borodinsky, 2015; Lee et al., 1997), but is later repressed through a Shh-Ca^2+^-PKA-CREB signaling axis that mediates Shh-dependent spinal cord neuron differentiation (Belgacem & Borodinsky, 2011, 2015).

PKA recruitment is not the only potential mediator of Hh non-canonical, Smo-dependent regeneration. The effects of inhibiting PKA on muscle regeneration are lower in magnitude compared with Smo inhibition, with the latter resulting in almost complete blockade of muscle replenishment in the regenerated tail. This may be the reason for epistasis when simultaneously inhibiting Smo and PKA pathways in regulating muscle regeneration. Alternatively, PKA may be one of multiple non-canonical Hh signaling pathways downstream of Smo acting on muscle regeneration (Figure 7).

The link between Smo and regeneration must lie in downstream transcription factors other than Gli, such as NFκB (Qu et al., 2013), MycN (Mani et al., 2020; Singh et al., 2018), CREB (Belgacem & Borodinsky, 2015) or JAK/STAT (Tian et al., 2015), which has previously been shown to regulate regeneration in *X*. *laevis* (Tapia et al., 2017). If specific signaling pathways downstream of Hh-dependent regeneration can be targeted, it may help limit the pleiotropic effects observed with direct manipulation of Smo (Wu et al., 2017).

Finally, PKA activity in spinal cord regeneration appears to act independently from Smo-dependent signaling. The effects of simultaneously inhibiting Smo and PKA are additive on spinal cord regeneration. The exact mechanisms by which PKA modulates regeneration, however, remain unclear. One possibility is that PKA may act as an inhibitor of Gli2, which would be in keeping with both our data showing Gli downregulation following amputation, and its developmental role as a repressor of canonical Hedgehog signaling (Hammerschmidt et al., 1996), in specific non-canonical, Ca^2+^ activity-dependent repression of Gli and activation of CREB during *X*. *laevis* spinal cord (Belgacem & Borodinsky, 2015).

Determining the exact mechanisms associated with Hh-dependent regeneration demands further investigation. One possibility comes from our discovery that stimulation of Smo induces a rapid increase in Ca^2+^ spikes in embryonic neurons in the developing spinal cord (Belgacem & Borodinsky, 2011) and this Ca^2+^ activity in combination with PKA causes a reversal in canonical Hh signaling, resulting in repression of Gli and activation of CREB transcriptional activity (Belgacem and Borodinsky, 2015). Since we have also found that Ca^2+^ activity is necessary for *Xenopus* larva tail and muscle regeneration following amputation (Tu and Borodinsky, 2014), our results showing an early reduction in Gli activity and activation of CREB, as well as the dependence of spinal cord and muscle regeneration on PKA activity, suggest that the aforementioned non-canonical pathway may be at least partially responsible for tissue regeneration.

Manipulation of Hh signaling has already shown great promise in treating a variety of conditions, including cancer (Ruat et al., 2014), neural injury and stroke (Bambakidis & Onwuzulike, 2012), cardiac ischemia (Dunaeva & Waltenberger, 2017), appendage regeneration (Chen et al., 2014; Singh et al., 2015), and even osteoporosis and obesity (Hadden, 2014). Based on the results from this study, we predict that selectively enhancing regeneration-specific, non-canonical Hh signaling in spinal cord and muscle during a critical period following injury might promote the repair and replenishment of functional tissues.

## Materials and Methods

### Animals

*Xenopus laevis* females were primed (50 units) and injected (350-400 units) with human chorionic gonadotropin to induce egg laying. Eggs were squeezed into 1X Marc’s Modified Ringer solution (MMR in mM: 110 NaCl, 2 KCl, 1 MgSO_4_, 2 CaCl_2_, 5 Hepes, 0.1 EDTA; pH to 7.8), fertilized with minced testis, and allowed to develop at a controlled temperature in 0.1X MMR. Mixed sex specimens were used at either 12-18 h (Nieuwkoop-Faber (NF) stage 11-16 embryos) or 60 h (NF stage 39-40 larvae) post fertilization.

### Tail amputation and pharmacological treatments

NF stage 39-40 larvae were anesthetized with 0.02% tricaine methanesulfonate (TMS, Syndel) with or without the specified drugs until non-responsive. Larvae were then amputated under a dissection stereoscope using a scalpel blade at approximately 1/3 of the length from the tail tip, where the tail begins to taper. The anesthetic was then washed out and the amputated larvae were incubated at 21-23°C in 10% MMR with vehicle or 10 μM cyclopamine (from 10-20 mM stock in DMSO; Sigma C4116), 1 μM SAG (from 5 mM stock in H_2_O; Calbiochem 566660), 10 μM GANT61 (from 10 mM stock in DMSO; Tocris 3191), and/or 10 μM KT5720 (from 10 mM stock in DMSO; Tocris 1288). For multi-day treatments, solutions were replaced daily.

### Whole mount immunostaining

Larvae were anesthetized in 0.02% TMS in 10% MMR, then fixed in 3.7% formaldehyde in 1X MEMFA saline (100 mM MOPS, 2 mM EGTA, 100 mM MgSO_4_) overnight at 4°C. Samples were bleached overnight in H_2_O_2_/Dent’s fixative, then permeabilized with 0.5% Triton X100 in 1X PBS (PBT) and blocked in 0.5% PBT + 2% BSA. Primary and secondary antibody incubations were performed in 0.1% PBT overnight at 4°C. Samples were washed in 0.5% PBT, mounted in 90% glycerol in 1X PBS and imaged within 1-4 days. Antibodies were obtained and used as follows: 1:300 Sox2 (R&D AF2018, RRID:AB_355110; neural stem cells), 1:100 12/101 (DSHB, RRID:AB_531892; skeletal muscle), 1:400 phospho-Serine10-histone-H3 (P-H3; Millipore 06-570, RRID:AB_310177, mitotic marker). All donkey secondary antibodies were used at 1:1500-1:2000 from Thermo Fisher: anti-Goat-Alexa-647 (A21447, RRID:AB_141844), anti-Goat-Alexa-594 (A11058, RRID:AB_2534105), anti-Mouse-Alexa-488 (A21202, RRID:AB_141607), anti-Rabbit-Alexa-594 (A21207, RRID:AB_141637), and anti-Rabbit-Alexa-647 (A31573, RRID:AB_2536183). Phospho-CREB (P-CREB) immunostaining required conditions as follows: Samples were fixed as above for 3 h at 4°C with gentle agitation, then washed in 0.1% PBT, dehydrated in methanol, and kept at -20°C overnight in methanol. Samples were then bleached at room temperature for 3 h, rehydrated, permeabilized as above, blocked in 10% BSA + 1.5% normal donkey serum, and placed in primary antibody solution: 1:400 Sox2, 1:100 12/101, and 1:1500 P-CREB (Cell Signaling 9198) for 4-5 days at 4°C, then treated as above for secondary antibody incubation, washes and mounting.

### Gli activity reporter

p8xGli-EGFP_CMV-iRFP670 was constructed as follows: The 8xGli-EGFP reporter plasmid was a gift from Prof. James Chen of Stanford University (Sasaki et al., 1997). The 8xGli-δcrystallin promoter was removed and spliced into the pXreg4-FireflyLuciferase_keratin-EGFP_CMV-RenillaLuciferase (courtesy of Dr. Yesser Hadj Balgacem, UC Davis, (Belgacem & Borodinsky, 2015) in place of the Xreg4-FL_keratin cassette to make p8xGli-EGFP_CMV-RenillaLuciferase. iRFP670 was then PCR-amplified from iRFP670-N1 ((Shcherbakova & Verkhusha, 2013), Addgene 45457) and swapped with Renilla Luciferase to make p8xGli-EGFP_CMV-RFP670. 150-200 pg p8xGli-GFP_CMV-iRFP670 were injected per embryo at the 2-4-cell stage. Individual cells were selected in Imaris (Bitplane) from greater than twice background intensity iRFP670+ cells and analyzed for mean EGFP and iRFP670 intensity. The construct was validated in 18 h post-fertilization (hpf), neural plate stage embryos +/-10 nM SAG. Imaging took place at 18 or 60-88 hpf embryos or larvae, respectively. Neural plate stage validation is presented as scatter plots, by iRFP670 (X) and EGFP (Y) signal intensity, with all iRFP670+ cells from all samples pooled (Figure 1-figure supplement 1). Post-amputation Gli activity is presented as two dimensional reconstructions of all iRFP670+ cells from all samples pooled, presented by each spot’s relationship to the amputation plane (x axis) and the spinal cord (y axis), with each iRFP670+ cell assigned a heat map intensity by EGFP:iRFP670 ratio (Figure 2).

### Gli2 knockdown

Morpholino antisense oligonucleotides targeted to block Gli2 translation ((Y. H. Belgacem & Borodinsky, 2015), 5’-GCACAGAACGCAGGTAATGCTCCAT-3’, Gli2-MO) and matched Gli2-PhotoMO (5’-ATGGAGCATTACPTGCGTTCT-3’) were ordered from Gene Tools. During and after injection, all steps were performed in the dark, using only >580 nm light for illumination. Embryos were injected at 4-cell stage with 4 nl of solution containing a total of 2 pmol each Gli2-MO and PhotoMO blocker with Cascade Blue-Dextran tracer and grown at 20-22°C until 32 hpf. Larvae were then split into two separate Petri dishes in 10% MMR, and either kept in the dark (inactive control) or exposed to 30 min of 365 nm UV transillumination, low output setting on a MaestroGen 240-V UV transilluminator (active MO). Larvae were then left at 20-22°C until 28 h post-UV, then amputated, and kept at 22-23°C until anesthesia, screening for tracer and fixation at 72 h post amputation (hpa).

### P-CREB+ cell quantification

Sox2 and P-CREB immunopositive cells within the spinal cord of immunostained whole mounts were separately analyzed on Imaris software (Bitplane). All cells were filtered for minimum 2X background Sox2 or P-CREB mean signal intensity, and background subtracted for mean Sox2 or P-CREB intensity. For P-CREB+ cells, 6 each of the dimmest acceptable and lowest intensity bright cells were visually selected for each sample, and their average values for P-CREB signal were used as threshold values for binning into dim (dimmest<P-CREB signal intensity<lowest bright) and bright (P-CREB signal intensity≥lowest bright) P-CREB-immunopositive cell sets. P-CREB+ cell counts are presented as the ratio of number of P-CREB+ cells in each category per total number of Sox2+ nuclei, for each 100-μm bin along the spinal cord posterior and anterior to the amputation plane.

### Live and fixed sample imaging

Live larvae anesthetized with 0.02% TMS in 0.1X MMR were imaged under a Nikon swept-field confocal microscope using 488 nm (EGFP) and 647 nm (iRFP670) lasers. Fixed samples were imaged on Nikon-A1 or C2 point laser-scanning confocal microscopes using 488 nm (Alexa488), 561 nm (Alexa594) and 640 nm (Alexa647) lasers.

### Western Blot assay

Nuclear and cytosolic fractions were obtained from stage 39-40 Gli2MO+PhotoMO larvae (three larvae for each group) to assess endogenous expression of Gli2. Briefly, larvae injected with Gli2MO + PhotoMO +/-UV (as described above) were frozen in liquid nitrogen, stored at -80°C, then homogenized in 25 mM Hepes pH 7.4, 50 mM NaCl, 2 mM EGTA, 5 mM MgCl_2_, protease inhibitors cocktail (784115, Thermo Fisher Scientific) on ice for 30 min and centrifuged for 10 min at 1000 g. Nuclear pellets and supernatant (cytosolic + membrane fractions) were resuspended in 2X protein loading buffer [125 mM Tris-HCl, pH 6.8, 4% SDS, 20% (w/v) glycerol, 0.005% Bromophenol Blue, 5% β-mercaptoethanol] and boiled for 5 min. Samples were run in 10% SDS-PAGE and transferred to PVDF membrane. PVDF membrane was probed with anti-Gli2 goat polyclonal (AF3635; RRID:AB_211902), 1:800 in 5% BSA at 4°C for 5 days, followed by incubation with horseradish peroxidase (HRP)-conjugated secondary antibody (711-035-152, Jackson ImmunoResearch; 1:10,000) and visualized by Western Lightning Plus-ECL, Enhanced Chemiluminescence Substrate (NEL103E001, Perkin Elmer). PVDF membranes were stripped in 0.2 M glycine HCl buffer, pH 2.5, 0.05% Tween for 20 min and re-probed with 1:500 anti-LaminII/III for nucleus-specific loading control (Cell Signaling 9087; RRID:AB_10896336) in 5% BSA.

### Experimental design and statistical analyses

All data were analyzed with Prism software (Graphpad). Data were first analyzed for normality, followed by parametric (normally distributed) or non-parametric tests (not normally distributed). When normally distributed and SDs were equal among groups, unpaired 2-tail t-test or ordinary one-way ANOVA followed by Tukey’s multiple comparisons test was used, when 2 or more groups were compared, respectively. When normally distributed and SDs were not equal among groups, Welch’s t-test or Brown-Forsythe and Welch ANOVA followed by Dunnett’s T3 multiple comparisons test was used, when 2 or more groups were compared, respectively. When data were not normally distributed, Mann-Whitney test or Kruskal-Wallis followed by Dunn’s multiple comparisons test was used, when 2 or more groups were compared, respectively. Graphed values are presented as a normalized percent of the experiment-matched control treatment, pooled across experiments. Significance was set to p<0.05. Number of samples and experiments are indicated in figure legends.

## Acknowledgements

We thank Dr. Yesser Hadj Belgacem for comments on the manuscript. This work was supported by NSF 1120796 and 1754340, NIH-NINDS R01NS073055, R01NS105886, R01NS113859 and Shriners Hospital for Children 86700-NCA grants to L.N.B. and Shriners Hospital for Children Postdoctoral Fellowship to A.M.H. We thank Christopher Hom for assistance with data acquisition and analysis and Olga Balashova and Jacqueline Levin for technical advice and assistance.

## Competing interests

The authors declare no competing financial or non-financial interests.

## Notes

### Competing Interest Statement

The authors have declared no competing interest.

