## Supplementary figures and images for "Non-canonical Hedgehog signaling regulates spinal cord and muscle regeneration"

### Supplemental Figures

#### Figure 2 Supplemental 1

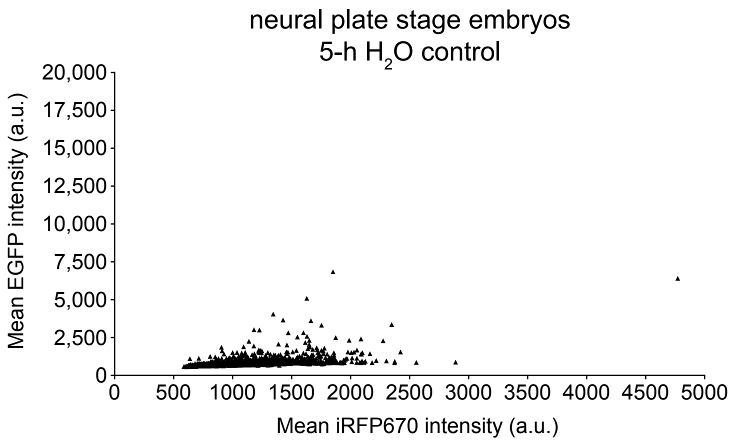


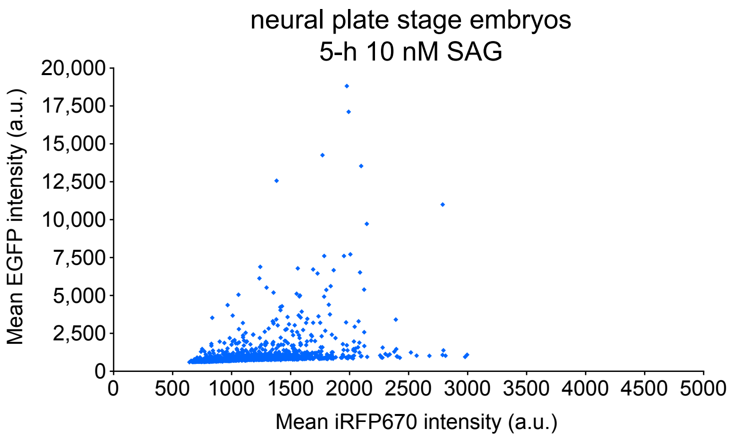

#### Figure 3 Supplemental 1

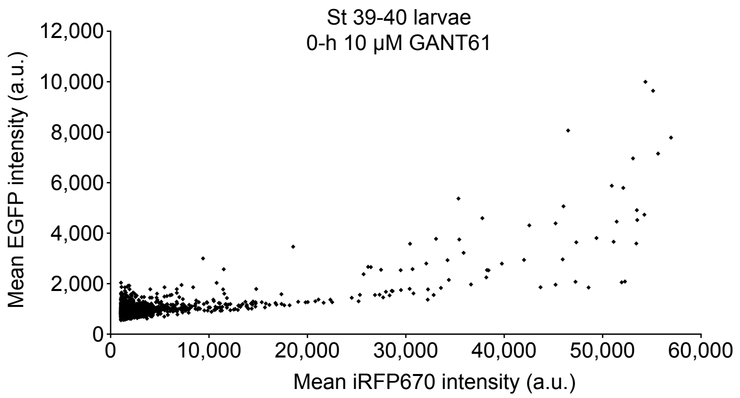


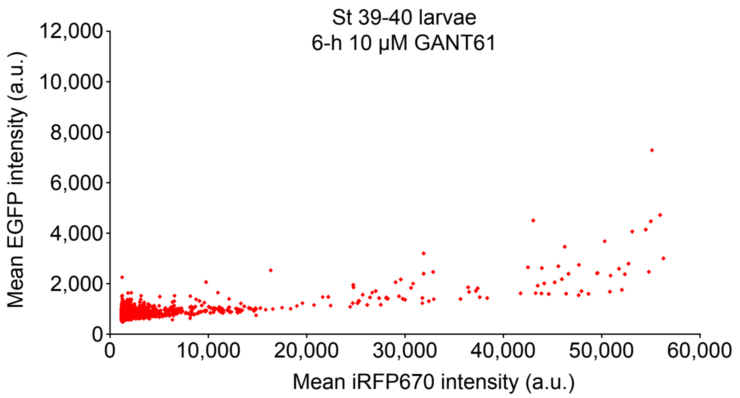


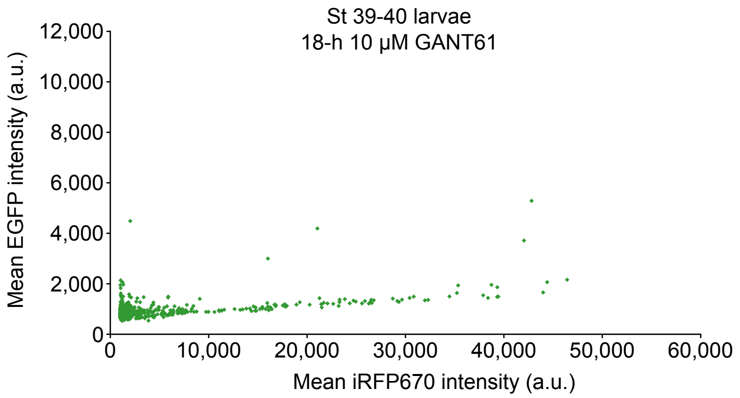

#### Figure 4 Supplemental 1

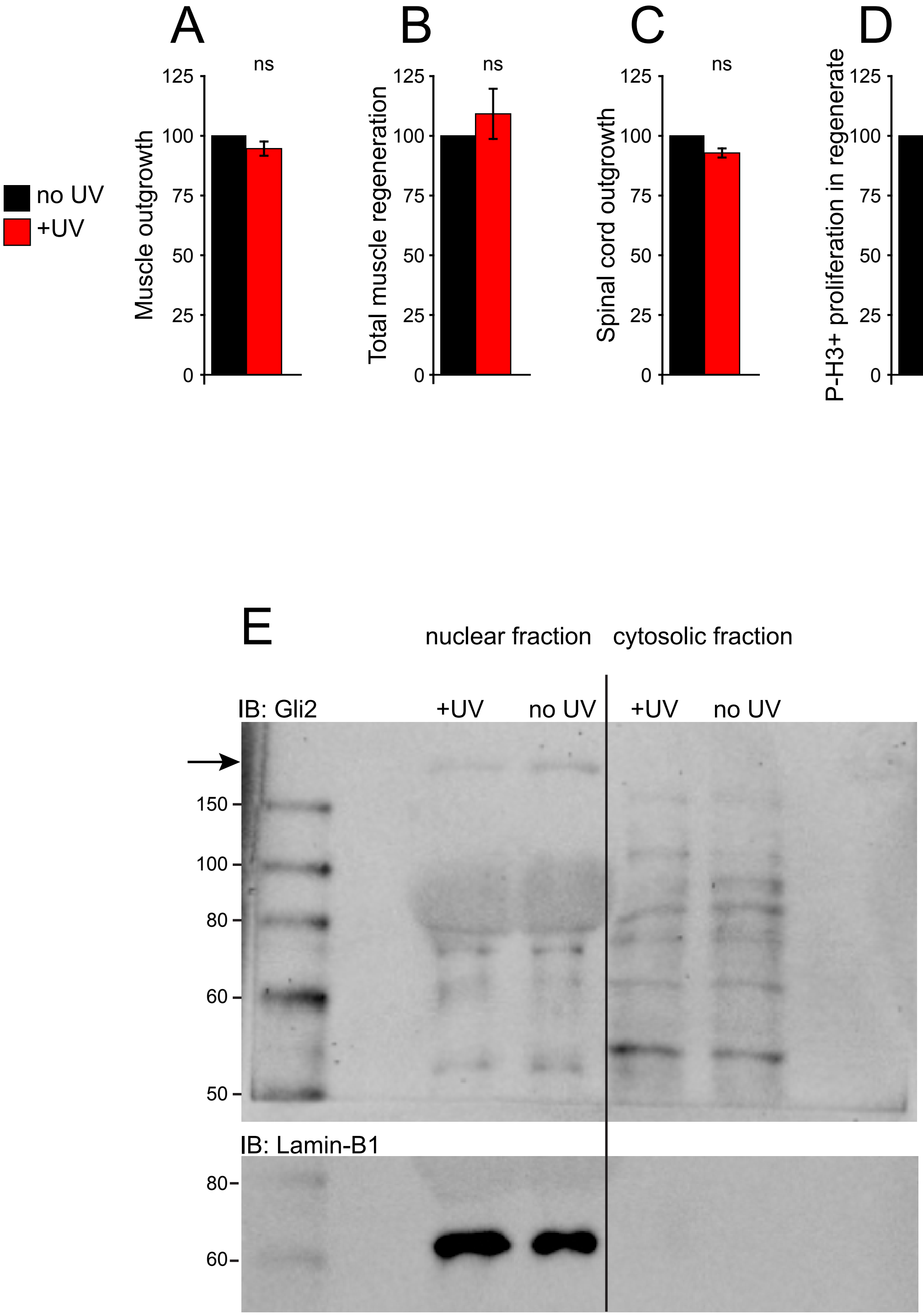

#### Figure 6 Supplemental 1

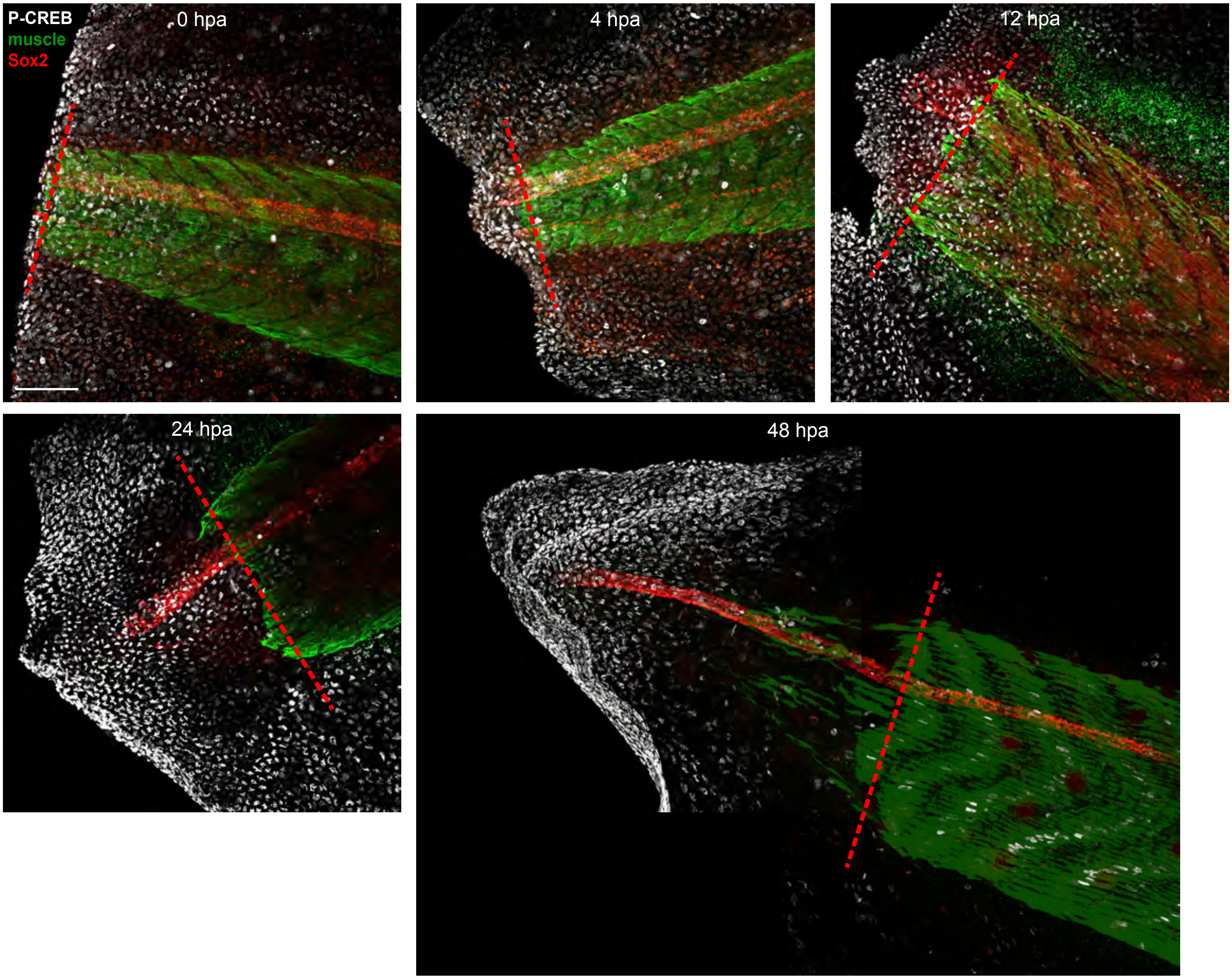
